# Inferring Sequence-Structure Preferences of RNA-Binding Proteins with Convolutional Residual Networks

**DOI:** 10.1101/418459

**Authors:** Peter K. Koo, Praveen Anand, Steffan B. Paul, Sean R. Eddy

**Author notes:** Corresponding Authors &.

## Abstract

To infer the sequence and RNA structure specificities of RNA-binding proteins (RBPs) from experiments that enrich for bound sequences, we introduce a convolutional residual network which we call ResidualBind. ResidualBind significantly outperforms previous methods on experimental data from many RBP families. We interrogate ResidualBind to identify what features it has learned from high-affinity sequences with saliency analysis along with 1st-order and 2nd-order *in silico* mutagenesis. We show that in addition to sequence motifs, ResidualBind learns a model that includes the number of motifs, their spacing, and both positive and negative effects of RNA structure context. Strikingly, ResidualBind learns RNA structure context, including detailed base-pairing relationships, directly from sequence data, which we confirm on synthetic data. ResidualBind is a powerful, flexible, and interpretable model that can uncover *cis*-recognition preferences across a broad spectrum of RBPs.

## Introduction

The life-cycle of RNA, including transcription, splicing, polyadenylation, transport, localization, and translation, is mediated by interactions with RNA-binding proteins (RBPs) (König *et al*, 2012). To gain mechanistic insight into RBP-regulated processes, it is essential to understand the specificity of RNA-RBP recognition. RNA-binding proteins contain modular binding domains that confer specific recognition preferences for RNA sequence and structure (Lunde *et al*, 2007; Campbell and Wickens, 2015; Jankowsky and Harris, 2015). To identify the RNA sequence preferences of an RBP, a variety of *in vitro* and *in vivo* experimental methods enrich for protein-bound RNA sequences (Lambert *et al*, 2014; Tome *et al*, 2014; Guenther *et al*, 2013; Ray *et al*, 2017; Licatalosi *et al*, 2008; Hafner *et al*, 2010; König *et al*, 2010; Sundararaman *et al*, 2016), and computational methods are used to deduce the consensus RNA sequence and/or structure features that these bound sequences share (Foat *et al*, 2006; Maticzka *et al*, 2014; Kazan *et al*, 2010; Maticzka *et al*, 2014; Orenstein *et al*, 2016; Alipanahi *et al*, 2015). Many analysis methods use position-weight-matrices (PWMs) or *k*-mers to model an RBP’s recognition code. Many of these methods make simplifying assumptions that do not necessarily capture all biologically important features, especially the positions of motifs and/or the influence of RNA secondary structure.

As powerful function approximators (Cybenko, 1989; Sonoda and Murata, 2015; Raghu *et al*, 2016; Shaham *et al*, 2016), deep neural networks (NNs) can learn a functional mapping between input genomic sequences and an output such as a bound/unbound label, or a binding affinity. Deep NNs do not necessarily have to make strong assumptions about what features of the input are important, because they autonomously build hierarchical representations of the data, in a machine learning approach called deep learning (LeCun *et al*, 2015). For supervised classification tasks, a deep NN is comprised of artificial neurons arranged in multiple layers: an input layer, one or more hidden layers, and an output layer where predictions are made (Goodfellow *et al*, 2016). Convolutional neural networks (CNNs) are a popular architectural choice that is well suited to detecting short contiguous patterns, such as motifs, along longer genomic sequences (Alipanahi *et al*, 2015; Zhou and Troyanskaya, 2015).

DeepBind was the first deep learning approach to analyze RBP-RNA interactions (Alipanahi *et al*, 2015). It demonstrated improved performance over PWM- and *k*-mer-based methods on the 2013-RNAcompete dataset, a standard benchmark dataset that consists of 244 *in vitro* affinity selection experiments that span across many RBP families (Ray *et al*, 2013). Recently, RCK (Orenstein *et al*, 2016), a *k*-mer-based approach, narrowly outperformed DeepBind. DeepBind’s design includes just a single convolutional layer which may limit its ability to learn feature representations that deeper CNNs with two or more convolutional layers could achieve. However, deeper models tend to be more difficult to train, and they can lack interpretability, making it challenging to understand what they have learned.

Recent advances in computer vision have introduced “residual networks” (He *et al*, 2016), a technique that makes it easier to train deep NNs, and an approach called “saliency analysis” that makes it easier to interpret what features they have learned (Simonyan *et al*, 2013; Springenberg *et al*, 2014; Shrikumar *et al*, 2016). Here we apply these advances to develop a convolutional residual network approach which we call ResidualBind.

## Results

### ResidualBind architecture

ResidualBind takes as input an RNA sequence and outputs a binding score prediction for a given RBP. ResidualBind can be decomposed into 5 stages (Fig. 1): (1) convolutional layer, (2) residual module, (3) mean-pooling layer, (4) convolutional layer, and (5) fully-connected output layer. Stage 1 is a convolutional layer that takes as input 41 nucleotide (nt) one-hot encoded sequences (**X**). 96 convolution filters, each of length 12 (**W**^(1)^), calculate a 1-dimensional cross-correlation across the sequence, outputting similarity scores at different positions, also known as feature maps (**Z**^(1)^). Each feature map is further processed with batch normalization (Ioffe and Szegedy, 2015) and a rectified linear unit (ReLU) activation (**H**^(1)^). In stage 2, the residual module takes **H**^(1)^ as input and performs two successive convolutional layers. **H**^(1)^ and the output of the 2 convolutional layers within the residual module are combined by an element-wise sum prior to a ReLU activation (**H**^(3)^). This skipped connection is the essence of the residual module, which foster training deeper networks by providing improved gradient flow (He *et al*, 2016). In stage 3, **H**^(3)^ is downsampled with a mean-pooling layer by averaging non-overlapping window sizes of 10 neurons separately for each filter, resulting in 3 neurons per filter that capture different spatial regions of the input sequence (**Ĥ** ^(3)^). Stage 4 employs a convolutional layer (**H**^(4)^) with 196 filters that have the same size as **Ĥ** ^(3)^, which is equivalently a fully-connected layer. Finally, stage 5 consists of a fully-connected output layer that takes **H**^(4)^ as input and performs a linear regression to a single output neuron (*O*), where binding score predictions are given.

**Figure 1:**
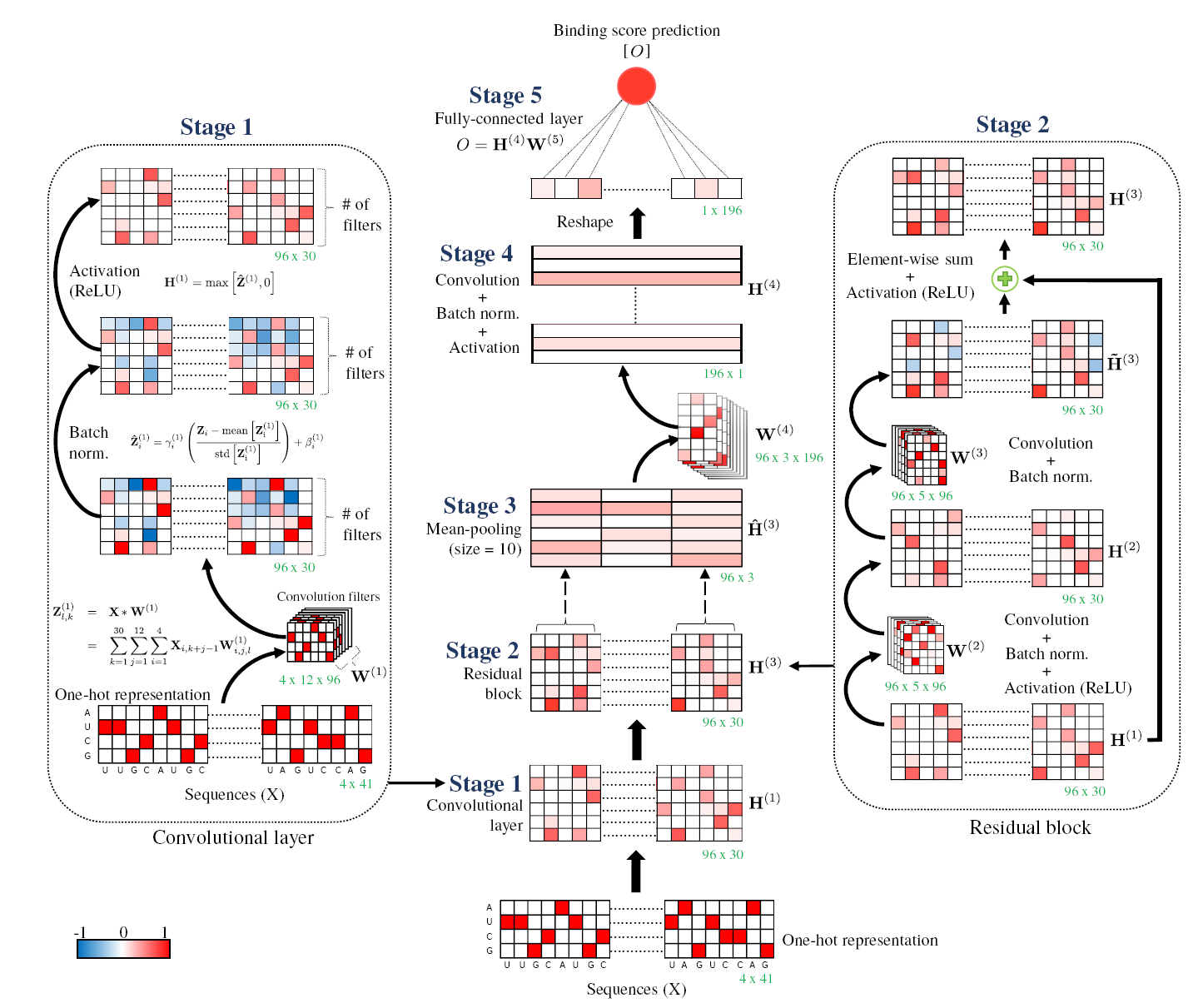
5-stage architecture of ResidualBind. ResidualBind stages consist of: (1) convolutional layer (details at left), (2) residual module (details at right), (3) mean-pooling layer, (4) convolutional layer, and a (5) fully-connected output layer, where the binding score predictions are given. The dimensions are shown in green on the bottom right of each feature map and filter. The convolutional layers in the residual module employ zero-padding to ensure the dimensions remain the same (not shown).

ResidualBind’s parameters, from each convolutional layer, batch normalization, and fully-connected layer, are determined through an iterative training procedure that minimizes the squared error between the predicted binding scores and the experimental binding scores (see Methods). We explored various hyperparameter settings by comparing the performance of candidate models with randomly-sampled hyperparamters on a validation dataset (Bergstra and Bengio, 2012). Many parameter settings perform comparably, so ResidualBind’s final hyperparameter setting was manually selected and fixed throughout all analyses in the paper.

### ResidualBind yields state-of-the-art predictions on the RNAcompete dataset

To benchmark the performance of ResidualBind against previously published methods, we used the 2013-RNAcompete dataset. RNAcompete consists of about 241,000 38-41 nt RNA sequences that are divided into two halves, ‘set A’ and ‘set B’, each of which consist of a different set of sequences that contain 8 copies of all possible 9-mers. RBPs are incubated with a 75-fold excess of RNA sequences so that the proportion of each bound RNA should scale with its relative affinity for the RBP at equilibrium. The experimental binding score given by RNAcompete is a normalized log-ratio of the fluorescence intensities of bound vs unbound RNA.

To compare ResidualBind against previous methods, we preprocessed experimental binding scores similar to (Orenstein *et al*, 2016; Alipanahi *et al*, 2015) by clipping large experimental binding scores to their 99.9th percentile value and normalizing to a *z*-score, a technique we refer to as clip-transformation. RNA sequences were converted to a one-hot representation with zero-padding added as needed to ensure all sequences had the same length of 41 nucleotides. For each RNAcompete experiment (i.e. each RBP) in the 2013-RNAcompete dataset, we trained a separate, randomly-initialized ResidualBind model on ‘set A’ sequences using experimental binding scores as training labels. Then, we predicted binding scores on ‘set B’ sequences using corresponding experimental binding scores to test the efficacy of our trained models.

We compared ResidualBind’s performance against MATRIXReduce (Foat *et al*, 2006), RNAcontext (Kazan *et al*, 2010), GraphProt (Maticzka *et al*, 2014), DeepBind (Alipanahi *et al*, 2015), and RCK (Orenstein *et al*, 2016). MATRIXReduce and RNAcontext are PWM-based methods. RNAcontext considers RNA secondary structure predictions along with RNA sequences. Graphprot employs a support vector machine with graph-kernel features. RCK is a method that replaces RNAcontext’s PWMs with *k*-mers. A common metric previously used to compare the performance of different models on the RNAcompete datasets is the Pearson correlation between model predictions and experimental binding scores on the held-out test set. Under this metric, ResidualBind significantly outperforms previously-reported methods, including the current state-of-the-art RCK (Fig. 2A). The mean Pearson correlation across all 244 experiments was 0.69 ± 0.17 for ResidualBind in comparison to 0.409 ± 0.166 for DeepBind and 0.460 ± 0.14 for RCK (errors are standard deviation of the mean). On a case-by-case basis, the Pearson correlations achieved by ResidualBind are nearly always found to be higher than RCK (Fig. 2B). This demonstrates that ResidualBind yields more accurate predictions that better reflect experimental binding scores in the RNAcompete dataset.

**Figure 2:**
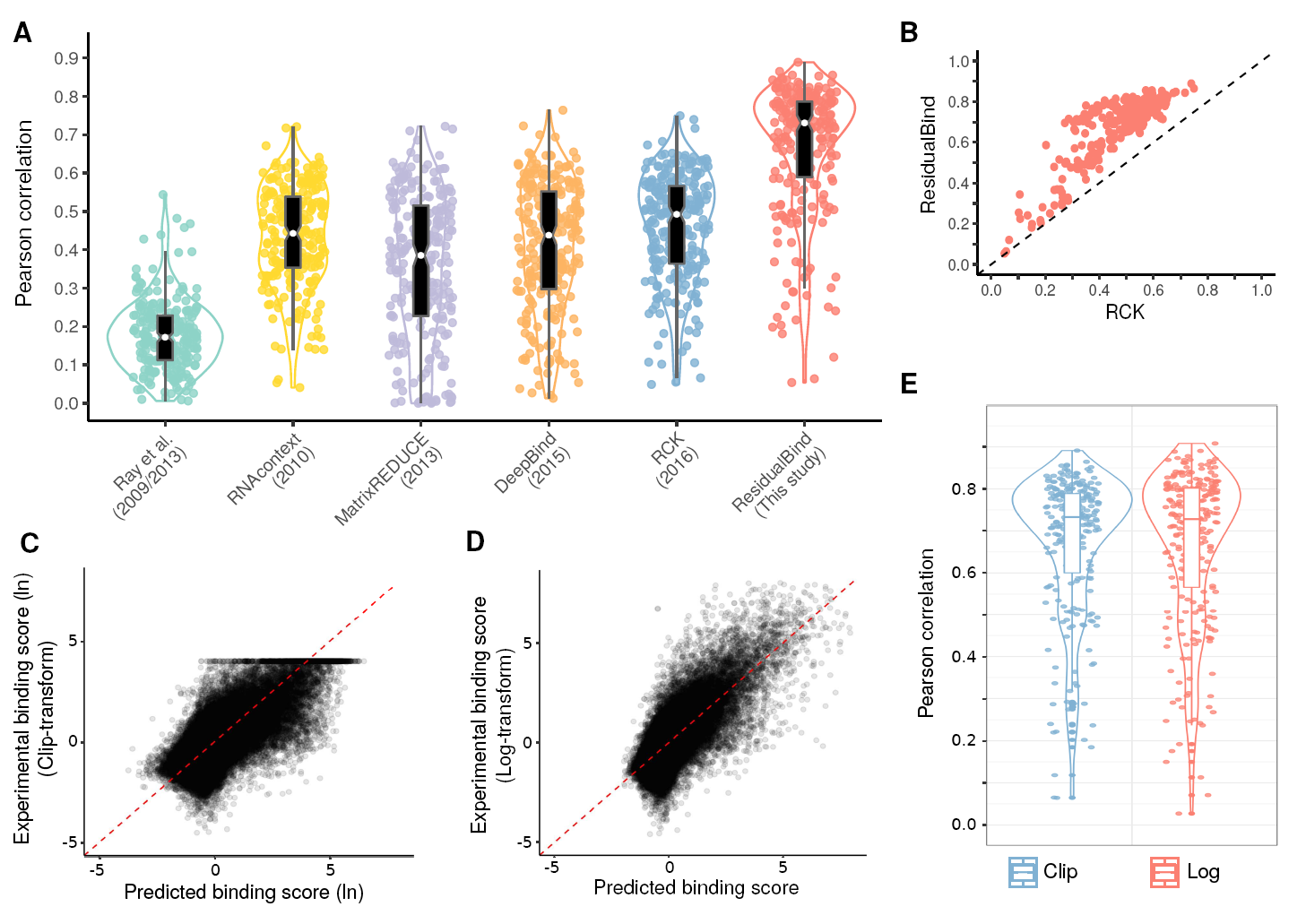
Performance comparison on the 2013-RNAcompete dataset. (A) Box-violin plot of Pearson correlations between experimental binding scores and predicted binding scores using different computational methods on the held-out test set for all 244 2013-RNAcompete experiments. Each data point represents one RBP experiment. Median value is shown as a white dot. (B) Scatter plot of ResidualBind’s Pearson correlations and RCK’s Pearson correlations. (C,D) Scatter plot of ResidualBind’s predicted binding scores and experimental binding scores from the test set of an RBP experiment in the 2013-RNAcompete dataset (RNCMPT00216) processed according to (C) clip-transformation and (D) log-transformation. (B-D) Red dashed line serves as a guide-to-the-eye for a perfect correlation. (E) Box-violin plot of the Pearson correlations between predicted binding scores given by ResidualBind and experimental binding scores of the test set for all 244 2013-RNAcompete experiments processed according to clip-transformation (blue) and log-transformation (red).

We noticed that clip-transformation adversely affects the fidelity of ResidualBind’s predictions for higher binding scores, the most biologically relevant regime (Fig. 2C). We prefer preprocessing experimental binding scores with a log-transformation similar to a Box-Cox transformation (see Methods), so that its distribution approaches a normal distribution while also maintaining their rank-order. With log-transformation, we found that ResidualBind yields higher quality predictions in the high-binding score regime (Fig. 2D), although average performance was essentially the same (Fig. 2E). Henceforth, our results will be based on preprocessing experimental binding scores with log-transformation.

### ResidualBind learns expected sequence motifs

We verified that ResidualBind learns expected consensus sequence motifs by performing *in silico* mutagenesis experiments, where we systematically make each nucleotide mutation to a canonical motif embedded in a synthetic sequence and ask how this alters ResidualBind’s binding score predictions. For each motif variant, we embed a single motif at the center of many different random RNA sequences, which serve to average out background noise. For example, Figure 3A shows results for RBFOX1, a well-studied protein whose canonical motif is ‘UGCAUG’ (dataset id: RNCMPT00168) (Auweter *et al*, 2006; Lambert *et al*, 2014; Lovci *et al*, 2013). ResidualBind finds that ‘GCAUG’ are the most important nucleotides in the RBFOX1 motif with minimal contributions from the 5’ U, which agrees with RBFOX1’s PWM in the CISBP-RNA database (ID: A2BP1 M159_0.6). The mean binding score for different variants correlate well with experimentally-determined ln *K*_*D*_ ratios of the variants and wild type measured by surface plasmon resonance experiments (Auweter *et al*, 2006) with a significant negative correlation of −0.763 (*p*-value= 0.046, *t*-test) (Fig. 3B).

**Figure 3:**
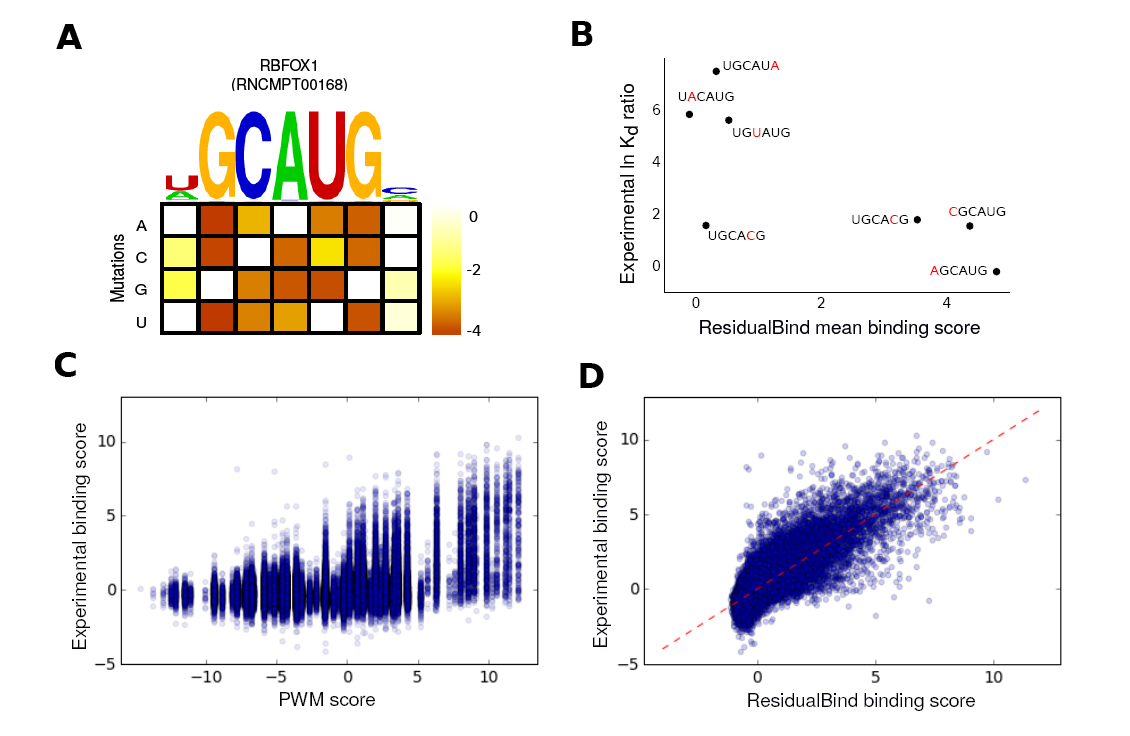
ResidualBind learns the expected RBFOX1 motif. (A) Heatmap of the difference between ResidualBind’s average binding score for 1,000 synthetic sequences embedded with each possible single nucleotide mutation of the canonical motif and the average binding score for 1,000 synthetic sequences embedded with a canonical RBFOX1 motif. A sequence logo of the RBFOX1 PWM from the CISBP-RNA database (ID: A2BP1 M159_0.6) is shown above. (B) Scatter plot of the experimental ln *K*_*d*_ ratio of the mutant to wild type measured via surface plasmon resonance (Auweter *et al*, 2006) versus ResidualBind’s average binding score for synthetic sequences embedded with a corresponding RBFOX1 variant. Each dot represents a different single nucleotide variant experiment, with the variant highlighted in red. (C,D) Scatter plot between experimental binding scores from the 2013-RNAcompete dataset (ID: RNCMPT00168) versus predicted binding scores given by (C) the maximum PWM score using an RBFOX1 PWM from the CISBP-RNA database, and (D) ResidualBind. (C,D) Each dot represents a different sequence in the test set.

Other methods, including DeepBind and RCK, learn a similar motif. Simply learning an RBP’s consensus motif is not sufficient to explain ResidualBind’s improved performance. For example, Figure 3C shows a scatter plot of the best PWM score for each test sequence versus experimental binding scores using an RBFOX1 PWM from the CISBP-RNA database (ID: A2BP1 M159_0.6). PWM scores do not correlate well with experimental binding scores (Pearson correlation=0.443 and Spearman rank correlation=0.315), particularly in the high binding score regime. In contrast, ResidualBind’s predicted scores better correlate with experimental binding scores (Fig. 3D, Pearson correlation 0.810, Spearman rank correlation=0.656), suggesting it is learning more complex features beyond a simple PWM.

### ResidualBind accounts for motif position, number and spacing

To identify nucleotides a deep network deems to be important, we employ saliency analysis (Shrikumar *et al*, 2016; Simonyan *et al*, 2013; Springenberg *et al*, 2014). Saliency analysis determines the sensitivity of predictions from perturbations to each nucleotide variant by calculating gradients of the outputs with respect to the inputs. We employ guided-backprop saliency analysis, which rectifies gradients through each ReLU activation to highlight the inputs that contribute toward increasing the prediction, a popular visualization technique in computer vision (Springenberg *et al*, 2014). The resultant “saliency map” can give insight into motifs that the neural network has learned and their spatial locations along a given sequence, without prior knowledge of the motif. Saliency analysis can only be applied on an individual sequence basis. For ease of viewing, we convert guided-backprop saliency maps into sequence logos (see Methods).

To show what ResidualBind has learned, we highlight saliency analysis for three sequences with high predicted binding scores that have a perfect match, a single mismatch, and two mismatches to the canonical RBFOX1 motif (Fig. 4A). We observe that a single intact RBFOX1 motif is sufficient for a high binding score (Fig. 4A, *i*), and sequences that contain mismatches to the canonical motif can also have high binding scores by containing additional ‘sub-optimal’ binding sites (Fig. 4A, *ii* -*iii*). This suggests that the number of motifs and possibly their spacing are relevant. This also raises the question of why a high RBFOX1 PWM hit doesn’t necessarily always correspond to high experimental binding scores (Fig. 3C). ResidualBind must be learning additional context that makes a high-scoring PWM hit a poor binding site. An example of such an effect could be a preference for the motif at certain positions.

**Figure 4:**
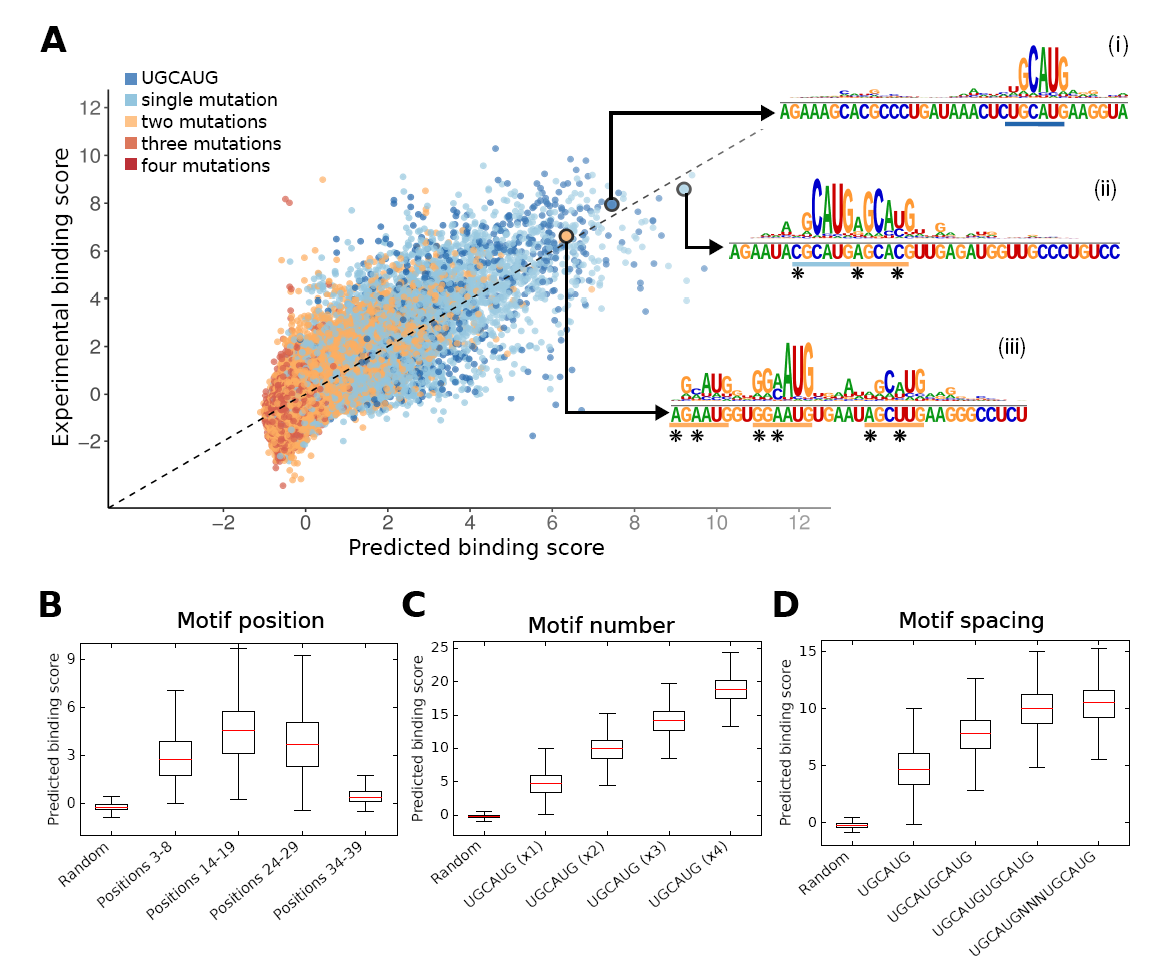
Features learned by ResidualBind for RBFOX1. (A) Scatter plot of experimental binding scores for test sequences in the 2013-RNACompete dataset for RBFOX1 versus ResidualBind’s predicted binding scores (Pearson correlation - 0.81). The color of each point is determined by the number of mutations between the canonical motif (UGCAUG) and its best match in the sequence. (i-iii) The inset shows sequence logos for saliency maps of select sequences with high predicted binding scores, a sequence with at best: (i) a perfect match, (ii) a single nucleotide mismatch, and (iii) a double nucleotide mismatch with the canonical RBFOX1 motif. The input RNA sequence is shown below each sequence logo. A colored bar below the sequence indicates the number of mutations with a star at locations of mismatches. (B) Box plot of binding score predictions for 1,000 synthetic sequences with an RBFOX1 motif embedded at various positions. (C) Box plot of binding score predictions for 1,000 synthetic sequences with varying numbers of embedded canonical RBFOX1 motifs in each sequence. (D) Box plot of binding score predictions for 1,000 synthetic sequences with two RBFOX1 motifs with varying degrees of separation.

We tested the effect of motif positional preference, motif numbers, and their spacings by systematically interrogating ResidualBind with synthetic sequences (see Methods). We observe that ResidualBind learns that RBFOX1 motifs near the 3’ end yield significantly lower binding scores compared to the rest of the sequence (Fig. 4B). This positional bias was confirmed by comparing the average experimental binding score for 5,000 sequences with the highest PWM scores, where the highest PWM hit is located within the final 3 positions of the 3’ end (0.036 ± 1.113, 1,062 total sequences) versus everywhere else (2.553 ± 2.306, 3,938 total sequences). This suggests that RBFOX1 has a nonspecific requirement of at least 3 nts flanking its specific motif for maximal binding. We also verified that ResidualBind learns the contribution of each motif is additive to the binding score (Fig. 4C), and the spacing between RBFOX1 motifs can decrease this effect when they are too close (Fig. 4D), which manifests biophysically through steric hindrance. These relationships are specific to RBFOX1 and differ in detail for other RBPs (Supplemental code).

### ResidualBind learns RNA secondary structures from sequence

In contrast to transcription factors, RNA structure is important for RBP recognition - some RBPs see structural targets, and some RBPs see unstructured targets that competing structures can block. Previous work, including RCK and RNAcontext (Orenstein *et al*, 2016; Kazan *et al*, 2010), has included RNA structure context prediction as an additional input with the sequence, and shown that this contributes to more accurate predictions. We explored whether similar inclusion of secondary structure predictions could improve ResidualBind’s performance.

The 2013-RNAcompete dataset was specifically designed to be weakly structured (Ray *et al*, 2013), so for these experiments we also used the 2009-RNAcomepete dataset, which consists of more structured RNA probes that include stem-loops for nine RBPs (Ray *et al*, 2009). Following procedures used in RCK and RNAcontext, we predicted two types of RNA secondary structure profiles for each sequence using RNAplfold (Lorenz *et al*, 2011) and a modified RNAplfold script (Kazan *et al*, 2010). RNAplfold yields secondary structure profiles that consist of the probability for each nucleotide to be either paired or unpaired (PU). The modified-RNAplfold script annotates each nucleotide with a predicted structural context of paired, hairpin-loop, internal loop, multi-loop, and external-loop (PHIME). Secondary structure profiles are incorporated into ResidualBind by creating additional input channels. The first convolutional layer now analyzes either 6 channels (4 channels for one-hot primary sequence and 2 channels for PU probabilities) or 9 channels (4 channels for one-hot primary sequence and 5 channels for PHIME probabilities).

Structure profiles do not increase ResidualBind’s performance (Fig. 5, A-B), but ResidualBind outper-forms RCK which does benefit from structure profiles (Fig. 5C). One possible explanation is that ResidualBind has already learned secondary structure effects from sequence alone. We compared the saliency representations learned by ResidualBind when trained on sequences with and without PU structural profiles for VTS1, a well-studied RBP whose SAM domain has a high affinity towards RNA hairpins containing ‘CNGG’ (Aviv *et al*, 2006b,a). The VTS1 motif in the CISBP-RNA database derived from analysis of the 2009-RNAcompete data is ‘GCUGG’. Saliency analysis shows that for ResidualBind trained on sequences alone, it has learned the canonical ‘(G)CNGG’ motif, but with other flanking nucleotides deemed important as well (Fig. 5D, *i* and *ii*). When PU secondary structural profiles are included as input, sequence-structure logos show that ResidualBind learns to recognize a VTS1 sequence motif in the context of a hairpin-loop structure.

**Figure 5:**
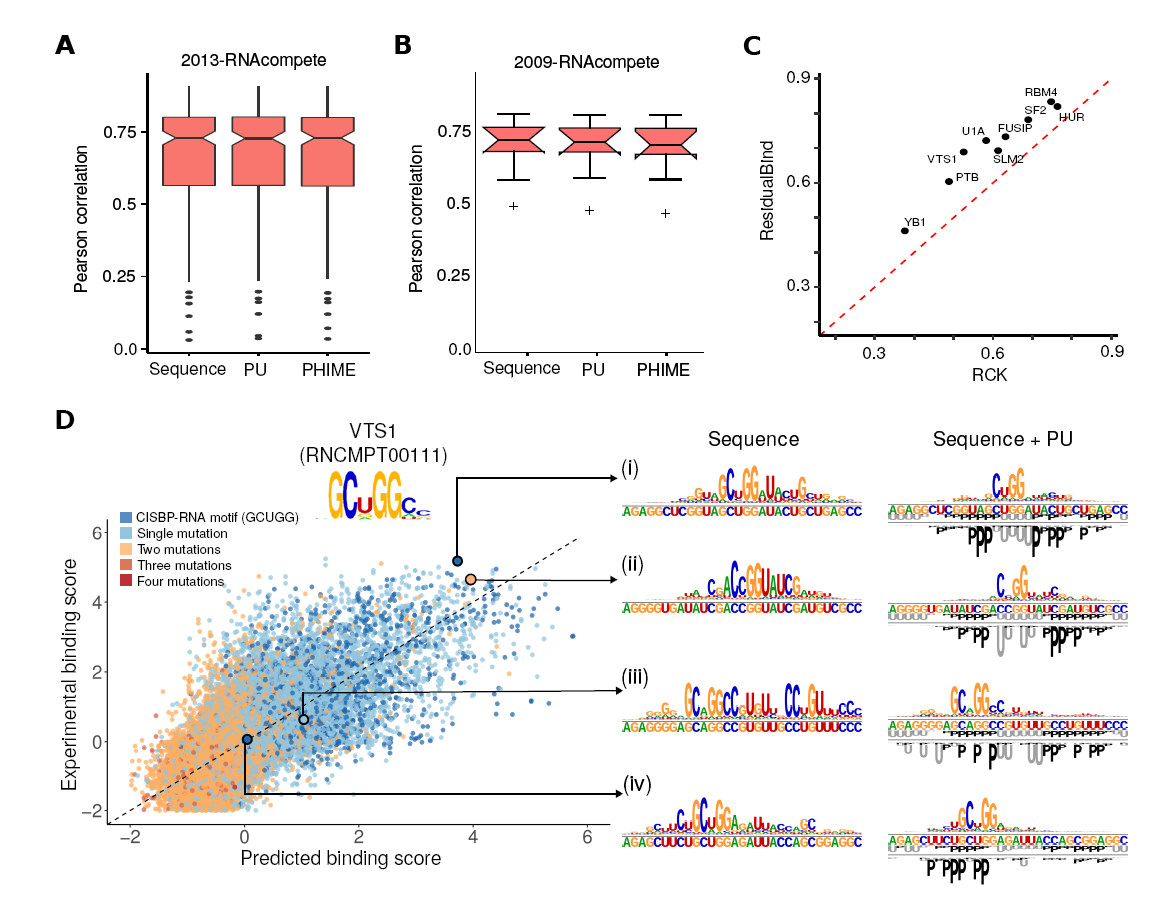
Effect of secondary structure. (A) Box plots of Pearson correlations between experimental and predicted binding scores for sequence, sequence+PU, or sequence+PHIME, for all 244 RBPs of the 2013-RNAcompete dataset. (B) Same, for 9 RBPs in the 2009-RNAcompete dataset. (C) Scatter plot of ResidualBind versus RCK of the Pearson correlation on test sets of 9 RBPs in the 2009-RNAcompete dataset. (D) Scatter plot of experimental binding scores for test sequences in the 2009-RNAcompete dataset for VTS1 versus ResidualBind’s predicted binding scores. The color of each point is determined by the number of mutations between the CISBP-RNA-derived motif (GCUGG) and the best match across the sequence. The inset shows sequence logos for saliency maps of representative sequences with high predicted binding scores (i-ii) and low predicted binding scores which contain the VTS1 motif (iii-iv). The sequence-structure logos contain the original sequence and structural profiles in the center, where ‘U’ represent unpaired (grey) and ‘P’ represents paired (black). The logo of the saliency map is placed on top for sequences and below for PU structural profiles.

We also looked at consensus CNGG motifs in unfavorable structural context. There are sequences with a VTS1 consensus CNGG sequence motif that have low experimental binding scores for which ResidualBind correctly predicts low binding scores (Fig. 5D). Saliency analysis on these low binding score sequences shows that ResidualBind trained on sequences alone identifies the canonical sequence motif and several flanking nucleotides as well (Fig. 5D, *iii* and *iv*). When ResidualBind is trained on sequences and PU structural profiles, saliency analysis of the same sequence shows that it recognizes the canonical sequence motif but in context of being overlapped by base pairs.

These results suggest that ResidualBind, trained on sequences alone, learns both positive and negative contributions of RNA structure context; when a structural profile is provided as input, it makes use of the profile instead, but either way gives similar performance. It seemed surprising to us that a CNN could learn detailed base pairing features, which requires the network to learn long-distance pairwise correlations (essentially XORs) on individual bases, using 1st layer feature detectors that are 12 nts wide. To test this, we designed a simple experiment where we trained ResidualBind to make a binary classification of whether or not an RNA sequence contains a hairpin structure from sequences only (see Methods). Briefly, we generated synthetic sequences with an 11 nt Watson-Crick paired stem and 7 nt loop (positive class); while background sequences contained random RNA sequences (negative class). After training, we gauge ResidualBind’s classification performance with the area under the receiver-operator-characteristic curve (AUC) on a withheld test set.

ResidualBind discriminated between hairpin sequences and unstructured sequences with an AUC of 0.9993. Saliency analysis on sequences with a high classification prediction for a hairpin loop shows that ResidualBind highlights nucleotides in the stem region of the synthetic RNA sequences (Fig. 6B). By itself, this evidence only suggests that ResidualBind may be recognizing long-distance complementary base pairing. To directly test whether it is, we employed a second-order *in silico* mutagenesis study, systematically scoring all 16 possible nucleotide pairs for every pair of positions (820 total pairs). If ResidualBind has learned detailed base-pairing interactions, we expect mutants with compensatory basepair substitutions to score well, while other score poorly. Figure 6C shows that this indeed is the result.

**Figure 6:**
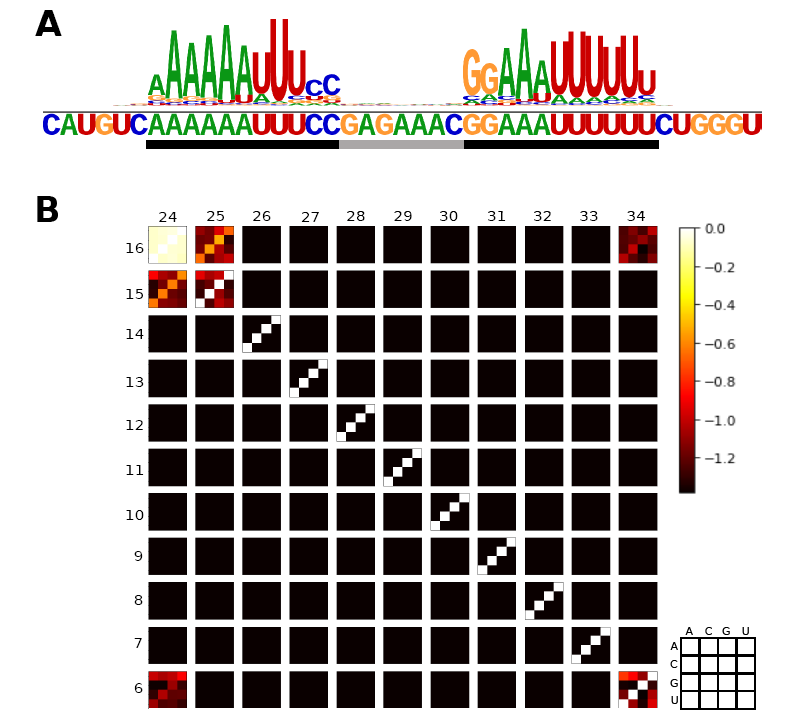
ResidualBind learns RNA base pairing from sequence input. (A) Saliency logo of a representative positive sequence, with stem and loop positions indicated by black and grey bars. (C) Heatmap of the difference between the average predicted binding scores of 2nd order mutagenesis sequences and the ‘wild type’ original sequences for all pairwise substitutions at stem positions 6-16 and 24-34 for 1,000 sequences with a high prediction for a hairpin loop. The lower-bound of the colorbar is clipped at −1.38, which corresponds to a 4 fold decrease in prediction from wild-type.

### ResidualBind learns other sequence biases

For many RBPs, the saliency map for top scoring sequences contains high GC content towards the 3’ end (Fig. 7). We did not observe any consistent secondary structure preference for the 3’ GC-bias. RBPs with similar binding motifs exhibit the same 3’ GC-bias trends; for instance, top scoring sequences for MSI (RNCMPT00040), Tb_0252 (RNCMPT00252) and RBM28 (RNCMPT00049), which all share a similar ‘GUAG’ motif, contain a 3’ GC-bias, but A2BP1 (RNCMPT00123) and ASD-1 (RNCMPT00180), which share a similar binding motif as RBFOX1, did not exhibit a noticeable 3’ GC-bias. We verified that ResidualBind had learned a 3’ GC-bias effect for a subset of RBPs by constructing and scoring synthetic sequences with and without an embedded motif and a 3’ GC bias (Fig. 7). We do not know the origin of this effect; it may be some sort of artifact that affects the abundance-levels of RNA probes. RNAcompete experiments determine the observed binding score as the ratio of the microarray fluorescence intensities from the affinity pull-down versus the remainder of unbound probes. This assumes that each RNA probe has approximately an equal abundance. However, if abundance levels are low, then the intensity ratio is more susceptible to statistical fluctuations. It is plausible that lower abundance probes tend to contain specific motifs. Many experimental steps in the RNAcompete protocol could lead to a GC-bias specifically on the 3’ end, including linker ligation, PCR amplification, transcription, among others (Wang *et al*, 2015; Sundararaman *et al*, 2016; Friedersdorf and Keene, 2014).

**Figure 7:**
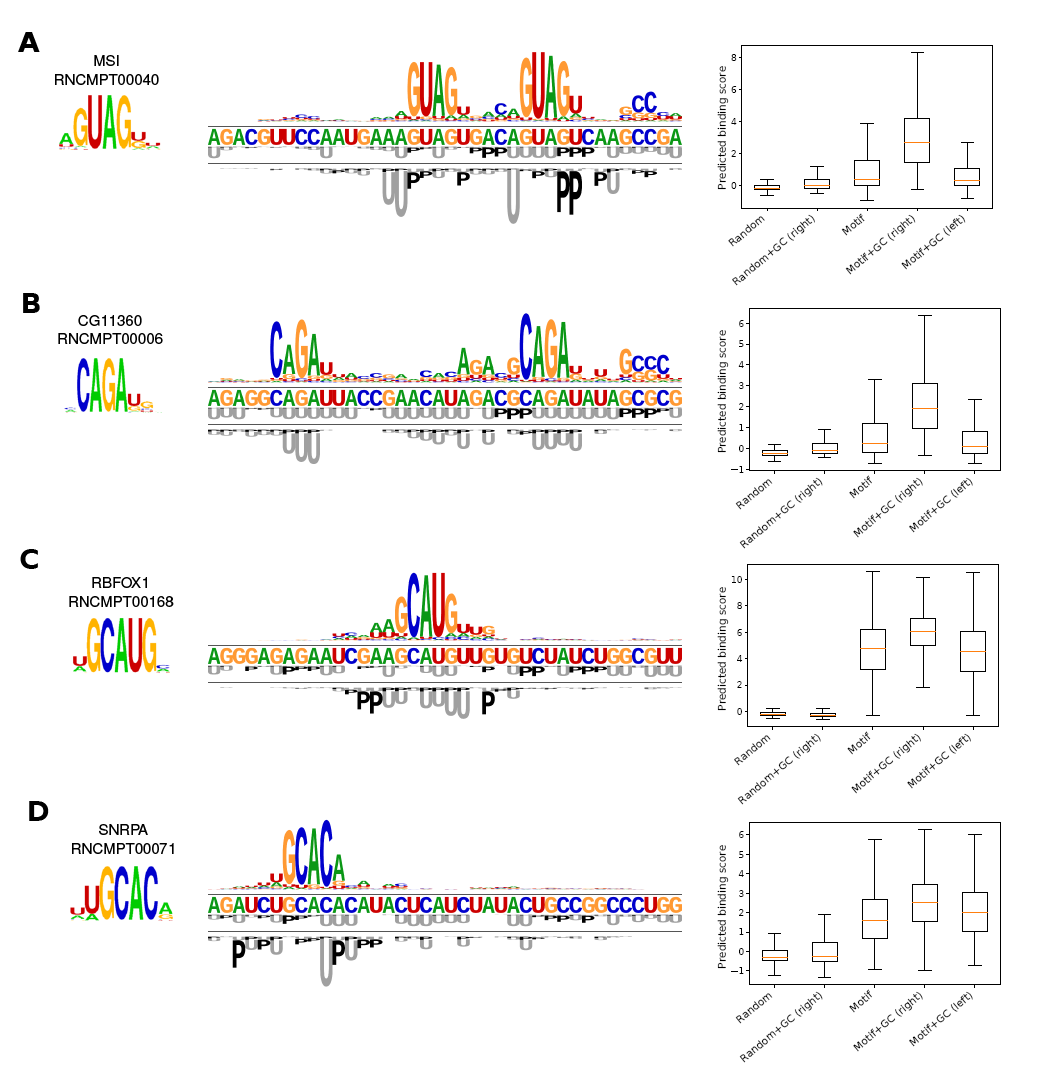
Effect of 3’ GC-bias. (A-D) CISBP-RNA motifs and representative sequence-structure logos of a high scoring sequence are shown for (A) MSI (RNCMPT00040), (B) CG11360 (RNCMPT00036), (C) RBFOX1 (RNCMPT00168), and (D) SNRPA (RNCMPT00071). Shown on the right is a box plot of the average binding scores predicted by ResidualBind for 1,000 sequences embedded with a pattern described in the x-label: ‘Random’ is random RNA sequences; ‘Random+GC (3’)’ is random RNA sequences with GCGCGC embedded on the 3’ end of the sequence; ‘Motif’ is random RNA sequences with a motif embedded at the center; ‘Motif+GC (3’)’ is random RNA sequences with a motif embedded at the center and GCGCGC embedded at the 3’ end of the sequence; ‘Motif+GC (5’)’ is random RNA sequences with a motif embedded at the center and GCGCGC embedded at the 5’ end of the sequence. The motif patterns used for MSI, CG11360, RBFOX1 and SNRPA were GUAG, CAGA, UGCAUG, UGCACA, respectively.

## Discussion

### ResidualBind’s design

ResidualBind is designed to autonomously learn discriminative features in RNA sequences that are predictive of RBP binding scores. The features it can learn are not limited to the convolutional filter size, because ResidualBind’s multiple convolutional layers can combine partial features learned in lower layers in a hierarchical manner (Koo and Eddy, 2018). ResidualBind employs a mean-pooling size that allows it to build representations across a large region, while maintaining general positions of individual features along the sequence. ResidualBind is a flexible model that can be broadly applied to a wide range of different RBPs without modifying hyperparameters for each specific experiment.

### lnterpretability

Tools for interpreting what deep learning models have learned are improving. We have shown that pertinent information learned by ResidualBind can be extracted via interrogation methods with saliency analysis and *in silico* experiments. While saliency analysis is a powerful approach to uncover features without any prior knowledge, it can only do so on an individual sequence basis. Thus, relevant features have to be deduced by observing general patterns across multiple sequences. Moreover, saliency methods yield seemingly noisy representations at times; in some cases, it is unclear whether certain nucleotides are important or not. To go a step further, we have found it useful to interrogate neural networks with synthetic sequences to uncover functional relationships between putative features and predictions. This approach allows us to directly test hypotheses of putative features with sequences in a controlled manner. Through saliency analysis and *in silico* experiments, we can see that ResidualBind has learned 3 *cis*-recognition principles that influence the sequence specificities of RBPs in the RNAcompete dataset: (1) multiple binding sites and their positions and spacings along a sequence, (2) the structural context and accessibility of each motif, and a (3) 3’ GC-bias. Using synthetic data is a powerful technique for model interpretability in regulatory geneomics. However, its application to natural language processing and computer vision may be limited, because it is not straightforward how to systematically synthesize data to test specific hypotheses in these other fields.

### *in vitro-to-in vivo* generalization

Ideally, a computational model trained on an *in vitro* dataset would learn principles that generalize to other datasets, including *in vivo* datasets. However, models trained on one dataset typically perform worse when tested on other datasets derived from different sequencing technologies/protocols (Weirauch *et al*, 2013). Different experimental protocols tend to produce biases that do not generalize (Wheeler *et al*, 2018; Wang *et al*, 2015; Sundararaman *et al*, 2016; Friedersdorf and Keene, 2014). For example, RNAcompete experiments: (1) perform a pull-down of a single binding domain of an RBP as opposed to the whole RBP, which is done *in ViVo*; (2) perform experiments under conditions that may not recapitulate *in vivo* conditions; (3) employ a limited set of short RNA probes that are designed for sequence content, *i.e.* all possible 9-mers, while *in vivo* sequences are biased through evolution; and (4) employ RNA probes that are designed to contain weak secondary structures, whereas *in vivo* sequences may contain more complicated structures (Mortimer *et al*, 2012). We found that ResidualBind learned a 3’ GC-bias in RNAcompete experiments that helps it to perform better than previous methods on test sequences derived from the same technology. However, we think the 3’ GC-bias is an example of a bias that arises from the particulars of the protocol. We do not think this feature is relevant to *in vivo* sequences, and our preliminary studies (of CLIP data) support this view. Better generalization to *in vivo* datasets is something that we hope to address in future work.

### Generality of ResidualBind

ResidualBind could be applied to understand signals in biological sequence data that enrich for any quantitative molecular sequence recognition phenotype, such as protein binding, histone modification, and chromatin accessibility, not just RNA binding. Our study demonstrates how a deep neural network like ResidualBind has a powerful ability to learn biological signals such as motifs, as well as confounding technical biases, and how such models can be interrogated with saliency analysis and synthetic data to reveal and test what principles the network has learned.

## Materials and methods

### RNAcompete dataset

#### Overview

We obtained the 2013-RNAcompete dataset from (Ray *et al*, 2013), where a full explanation of the data can be found. Briefly, the 2013-RNAcompete experiments consist of an Agilent 244K microarray which contains 240,000 oligonucleotides of length 30-41 nucleotides. The library of oligo sequences was designed to ensure that all possible combinations of 9-mers are sampled at least 16 times. The probes are divided into two sets - ‘set A’ (120,326 sequences) and ‘set B’ (121,031 sequences). A pool of RNA sequences is prepared from the microarray and incubated with a recombinantly-expressed RBP of interest tagged with glutathione S-transferase (GST). The total RNA concentration is in 75-fold molar excess over protein (20 nM protein, 1,500 nM RNA) so that at equilibrium the proportion of each sequence bound to RBP is expected to be proportional to its affinity. Bound RNA is recovered by a pull down, labeled with Cy5, and combined with the Cy3-labeled input RNA pool. The mixture of labelled RNA is hybridized to a copy of the same Agilent 244K microarray. The provided binding score for each sequence is the log-ratio of the fluorescence intensities of pull-down versus input, which serves as a measure of binding affinity and therefore sequence preference, that is normalized to remove various technical biases. The 2013-RNAcompete dataset consists of 244 experiments for 207 RBPs using only weakly structured probes (Ray *et al*, 2013). We also obtained the 2009-RNAcompete dataset, which is comprised of nine RBP experiments that employ a mix of probes that were predicted to be either weakly structured or contain stem-loops (Ray *et al*, 2009).

#### Preparation of RNAcompete datasets

Each sequence from ‘set A’ and ‘set B’ was converted to a one-hot representation. The one-hot sequences and the experimental binding scores for each experiment in the 2013-RNAcompete dataset were stored in a single HDF5 file (availability: eddylab.org/publications/ KooEddy19/residualbind_data.tar.gz). We filtered out sequences with a binding score of NaN. We then performed either clip-transformation or log-transformation. Clip-transformation is performed by clipping the extreme binding scores to the 99.9th percentile binding score. Log-transformation processes the binding scores according to the function: log (*S –S*^*MIN*^ + 1), where *S* is the raw binding score and *S*^*MIN*^ is the minimum value across all raw binding scores. This monotonically reduces extreme binding scores while maintaining their rank order, and also yields a distribution that is closer to a Normal distribution. The processed binding scores of either clip-transformation or log-transformation were converted to a z-score. We randomly split set A sequences to fractions 0.9 and 0.1 for the training set and validation set, respectively. Set B data was held out and used for testing.

#### Secondary structure predictions

For each sequence, structural profiles, which consist of predicted paired-unpaired (PU) probabilities for each nucleotide, were calculated using RNAplfold (Bernhart *et al*, 2005). Structural profiles consisting of predicted paired probabilities of five types of RNA structure - paired, hairpin-loop, internal loop, multi-loop, and external loop (PHIME) - were calculated using a modified RNAplfold script (Kazan *et al*, 2010). For each sequence, the window length (-W parameter) and the maximum spanning base-pair distance (-L parameter) were set to the full length of the sequence.

### *In silica* experimental design

#### RBFOX1 mutagenesis

1,000 random RNA sequences, each 41 nucleotides long, were simulated from an equiprobable i.i.d. uniform sequence model. A canonical RBFOX1 motif, *i.e.* UGCAUG, or each of the 21 possible single-nucleotide variants of it was embedded in each random sequence at position 22-28, resulting in 22,000 total sequences. Since the same 1,000 random sequences to embed each motif variant, the same background noise distribution is shared across different variants.

#### Binding site location

1,000 random RNA sequences, each 41 nucleotides long, were simulated from a uniform sequence model. 4,000 additional sequences were generated by embedding each of the 1,000 sequences with a single RBFOX1 motif at positions 4-9, 14-19, 24-29, and 34-39.

#### Multiple binding sites

1,000 random RNA sequences, each 41 nucleotides long, were simulated from a uniform sequence model. 4,000 additional sequences were generated by embedding each of the 1,000 sequences with: 1 RBFOX1 motif (positions: 18-23), 2 RBFOX1 motifs (positions: 11-16 and 18-23), 3 RBFOX1 motifs (positions: 11-16, 18-23, and 26-31), and 4 RBFOX1 motifs (positions: 4-9, 11-16, 18-23, and 26-31).

#### Motif separation

1,000 random RNA sequences, each 41 nucleotides long, were simulated from a uniform sequence model. 4,000 additional sequences were created by embedding 4 patterns starting at position 16 in each of the 1,000 sequences: UGCAUG, UGCAUGAUG, UGCAUGUGCAUG, and UGCAUGNNNUG-CAUG, where N represents a random nucleotide.

#### GC-bias

1,000 random RNA sequences, each 41 nucleotides long, were simulated from a uniform sequence model. 4,000 additional sequences were created by embedding one of 4 patterns: (1) GCGCGC at the 3’ end at positions 35-40, (2) a motif embedded at the center (starting at position 17), (3) a motif embedded at the center and GCGCGC at the 3’ end, and (4) a motif embedded at the center and GCGCGC at the 5’ end at positions 1-6. The embedded motifs were GUAG (MSI, RNCMP00040), CAGA (CG11360, RNCMP00006), UGCAUG (RBFOX1, RNCMPT00168), and UGCAC (SNRPA, RNCMPT00071).

#### RNA hairpin

Synthetic 41 nt RNA stem-loop sequences were generated as random sequences of equiprobable composition, with positions 25-35 fixed to be complementary to positions 7-17. Unstructured synthetic sequences were generated as random sequences of equiprobable composition. 100,000 synthetic RNA stem-loop sequences (positive class) and 100,000 random RNA sequences were randomly split 80% to a training set, 10% validation set, and 20% test set.

### Saliency analysis

Saliency analysis was performed by guided-backprop which calculates the gradients of the outputs with respect to the inputs (Springenberg *et al*, 2014). In contrast to standard backpropagation, guided-backprop rectifies negative gradients that pass through each ReLU activation in each hidden layer.

To generate sequence logos, we normalized the guided-backprop-generated saliency maps by dividing the maximum absolute value across the saliency map. Next, we applied an exponential filter according to: 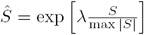, where *S* is the saliency map, and λ is a scaling factor that we set to 3 in this paper. This filtering step supresses small gradient signals and enhances larger gradient signals. We have previously found this filtering step to empirically yield sequence logos that better reflect PWMs from analysis on synthetic sequences (Koo and Eddy, 2018). We then separately normalized each nucleotide position by dividing by the sum of the filtered saliency map, thereby providing a probability for each nucleotide at each position. To generate a sequence logo, the normalized saliency value of each nucleotide *a* at position *i* was scaled according to: *Ŝ*_*a,i*_ × *I*_*i*_, where *I*_*i*_ = 2 + ∑ _a_ *Ŝ _a,i_* log_*2*_ *Ŝ _a,i_*.

### ResidualBind

#### Training ResidualBind

We trained a separate ResidualBind model on sequences in ‘set A’ for each RNAcompete experiment by minimizing the mean squared-error loss function between the model predictions and the experimental binding scores. All models were trained with mini-batch stochastic gradient descent (mini-batch of 100 sequences) with Adam updates using recommended default parameters with a constant learning rate of 0.0003 (Kingma and Ba, 2014). Dropout was applied after each convolutional layer with a dropout probability of 0.2, 0.1, 0.2, 0.5, sequentially from layer 1 to layer 4. During training, we also employed L_2_-regularization with a strength equal to 10^*-*6^. Training was stopped when the loss on the validation dataset does not improve for 20 epochs (early stopping). Optimal parameters were selected by the epoch which yields the lowest loss on the validation dataset. The parameters of each model were initialized according to He initialization (He *et al*, 2015). Training was performed on a NVIDIA GTX Titan X Pascal graphical processing unit with acceleration provided by cuDNN libraries (Chetlur *et al*, 2014). On average, training was complete after about 50 epochs, where each epoch takes 1-3 seconds on average.

#### Modified-ResidualBind for binary classification

To make binary classification predictions as opposed to a continuous binding score prediction, we modified ResidualBind by placing its linear output through a logisitic function, also known as a sigmoid activation. The mean-squared loss function was also changed to a binary cross-entropy loss function using binary labels corresponding to the presence or absence of a RNA hairpin structure in the sequence.

### NeuralBinder

Our implementation of ResidualBind utilizes a custom-written, open-source Python package that we call NeuralBinder. NeuralBinder includes high-level Python scripts and Python notebooks that provide step-by-step instructions on how to build, optimize, save, and load neural network models, how to evaluate a trained model on new sequences, and how to perform saliency analysis. NeuralBinder in turn wraps Deepomics, a custom-written module that contains high-level APIs written on top of TensorFlow (Abadi *et al*, 2016) to build, train, test, and evaluate neural network models. NeuralBinder can be applied to perform similar analyses on any affinity selection-based experiments, such as RNAcompete, SELEX (Tuerk and Gold, 1990), ChIP-seq, or CLIP-seq. Code to replicate this study is provided in the supplementary materials. An updated version of NeuralBinder that is continually under development can be found via: https://github.com/p-koo/neuralbinder.

## Acknowledgements

We thank Harleen Saini, Tom Jones, Tim Dunn, Elena Rivas, Fred Davis, Nick Carter, and other members of the Eddy/Rivas lab for helpful discussions and feedback.

